# Greengenes2 enables a shared data universe for microbiome studies

**DOI:** 10.1101/2022.12.19.520774

**Authors:** Daniel McDonald, Yueyu Jiang, Metin Balaban, Kalen Cantrell, Qiyun Zhu, Antonio Gonzalez, James T. Morton, Giorgia Nicolaou, Donovan H. Parks, Søren Karst, Mads Albertsen, Philip Hugenholtz, Todd DeSantis, Se Jin Song, Andrew Bartko, Aki S. Havulinna, Pekka Jousilahti, Susan Cheng, Mike Inouye, Teemu Niiranen, Mohit Jain, Veikko Salomaa, Leo Lahti, Siavash Mirarab, Rob Knight

**Affiliations:** Department of Pediatrics, UC San Diego School of Medicine, La Jolla, CA, USA; Department of Electrical and Computer Engineering, UC San Diego, La Jolla, CA, USA; Bioinformatics and System Biology Program, UC San Diego, La Jolla, CA, USA; Department of Computer Science and Engineering, UC San Diego, La Jolla, CA, USA; School of Life Sciences, Arizona State University, Tempe, AZ, USA; Biodesign Center for Fundamental and Applied Microbiomics, Arizona State University, Tempe, AZ, USA; Biostatistics & Bioinformatics Branch, Eunice Kennedy Shriver National Institute of Child Health and Human Development, National Institutes of Health, Bethesda, MD, USA; Halicioglu Data Science Institute, UC San Diego, La Jolla, CA, USA; Australian Centre for Ecogenomics, School of Chemistry and Molecular Biosciences, The University of Queensland, St Lucia, QLD 4072, Australia; Department of Obstetrics and Gynecology, Columbia University, New York, NY, USA; Department of Chemistry and Bioscience, Aalborg University, Aalborg, Denmark; Department of Informatics, Second Genome, Brisbane, CA, USA; Center for Microbiome Innovation, Jacobs School of Engineering, UC San Diego, La Jolla, CA, USA; Finnish Institute for Health and Welfare, Helsinki, Finland; Institute for Molecular Medicine Finland, FIMM-HiLIFE, Helsinki, Finland; Division of Cardiology, Brigham and Women’s Hospital, Boston, MA, USA; Cedars-Sinai Medical Center, Los Angeles, CA, USA; Cambridge Baker Systems Genomics Initiative, Baker Heart and Diabetes Institute, Melbourne, VIC, Australia; Cambridge Baker Systems Genomics Initiative, Department of Public Health and Primary Care, University of Cambridge, Cambridge, UK; Division of Medicine, Turku University Hospital and University of Turku, Turku, Finland; Sapient Bioanalytics LLC, San Diego, CA, USA; Department of Computing, University of Turku, Turku, Finland; Department of Bioengineering, UC San Diego, La Jolla, CA, USA

## Abstract

16S rRNA and shotgun metagenomics studies typically yield different results, usually attributed to biases in PCR amplification of 16S rRNA genes. Here, we introduce Greengenes2 and show that differences in reference phylogeny are more important. By inserting sequences into a whole-genome phylogeny, we show that 16S rRNA and shotgun metagenomic data generated from the same samples agree in principal coordinates space, taxonomy, and in phenotype effect size when analyzed with the same tree.

## Body

Shotgun metagenomics and 16S rRNA gene amplicon (16S) studies are widely used in microbiome research, but investigators using different methods typically find their results hard to reconcile. This lack of standardization across methods limits the utility of the microbiome for reproducible biomarker discovery.

A key problem is that whole-genome resources and rRNA resources depend on different taxonomies and phylogenies. For example, Web of Life (WoL) ^1^ and the Genome Taxonomy Database (GTDB) ^2^ provide whole-genome trees that cover only a small fraction of known bacteria and archaea, while SILVA ^3^ and Greengenes ^4^ are more comprehensive but most often not linked to genome records.

We reasoned that an iterative approach could yield a single massive reference tree that unifies these different data layers (e.g., genome and 16S rRNA records). We began with a whole-genome catalog of 15,953 bacterial and archeal genomes evenly sampled from NCBI, and reconstructed an accurate phylogenomic tree by summarizing evolutionary trajectories of 380 global marker genes using the new workflow uDance. This work, namely Web of Life version 2 (WoL2), represents a significant upgrade from the previously released WoL1 (10,575 genomes) ^15^. Then, we added 18,356 full-length 16S rRNA sequences from the Living Tree Project January 2022 release ^6^, 1,725,274 near-complete 16S rRNA genes from Karst et al. ^7^ and the EMP500 ^8^, and all full-length 16S rRNA sequences from GTDB r207 to the genome-based backbone with uDance v1.1.0, producing a genome supported phylogeny with 16S rRNA explicitly represented. Finally, we inserted 23,113,447 short V4 16S rRNA Deblur v1.1.0 ^9^ amplicon sequence variants (ASVs) from Qiita (retrieved Dec. 14, 2021) ^10^ as well as mitochondria and chloroplast 16S from SILVA v138 using DEPP v0.3^11^. This final step represents ASVs from over 300,000 public and private samples in Qiita including the entirety of the Earth Microbiome Project ^12^ and American Gut Project/Microsetta ^13^ (Fig. 1A). Our use of uDance ensured the genome-based relationships are kept fixed and relationships between full-length 16S rRNA sequences are inferred. For short fragments, we kept genome and full-length relationships fixed and inserted fragments independently from each other. Following deduplication and quality control on fragment placement, this yielded a tree covering 21,074,442 sequences from 31 different Earth Microbiome Project Ontology (EMPO) EMPO_3 environments, of which 46.5% of species-level leaves were covered by a complete genome. Taxonomic labels were decorated onto the phylogeny using tax2tree v1.1 ^4^. The input taxonomy for decoration used GTDB r207, combined with the Living Tree Project January 2022 release. Taxonomy was harmonized prioritizing GTDB including preserving the polyphyletic labelings of GTDB (see also Online Methods). Taxonomy will be updated every six months using the latest versions of GTDB and LTP.

**Figure 1.**
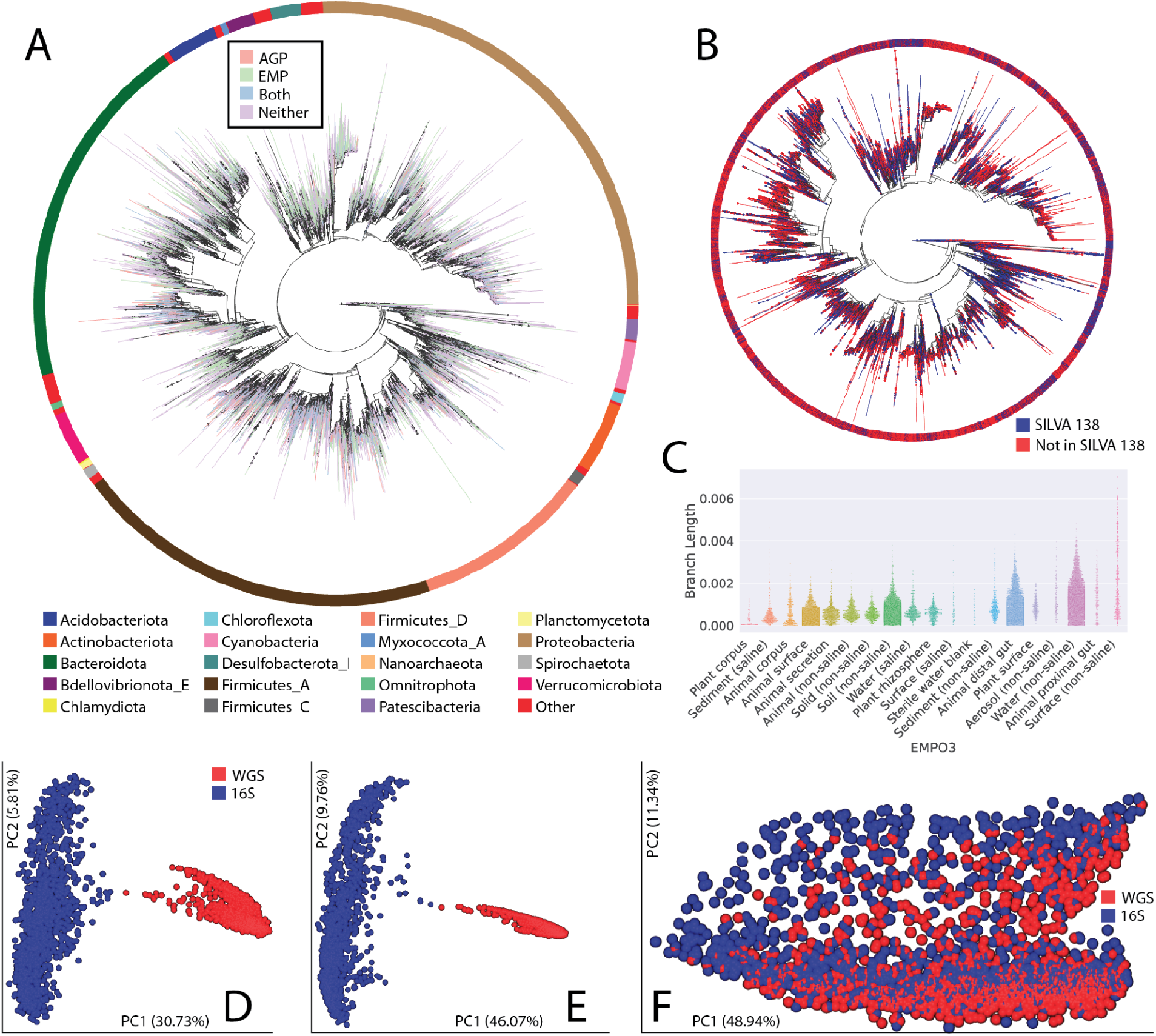
(A) The Greengenes2 phylogeny rendered using Empress ^18^ with amplicon sequence variant multifurcations collapsed, tip color indicating representation in the American Gut Project (AGP), the Earth Microbiome Project (EMP), both or neither, and with the top 20 represented phyla depicted in the outer bar. (B) The same collapsed phylogeny, colored by the presence or absence of a best BLAST ^19^ hit from SILVA 138.The bar depicts the same coloring as the tips. (C) Earth Microbiome Project samples and the amount of novel branch length, normalized by the total backbone branch length, added to the tree through amplicon sequence variant fragment placement. Note that sample counts are not even across EMPO3 categories. (D) Bray-Curtis applied to paired 16S V4 rRNA amplicon sequence variants and whole genome shotgun samples from The Human Diet Microbiome Initiative subset of The Microsetta Initiative. (E) Same data as (D) but computing Bray-Curtis on genus collapsed data. (F) Same data as (D-E) but using weighted UniFrac at the ASV and genome identifier level.

Greengenes2 is much larger than past resources in its phylogenetic coverage, as compared to SILVA (Fig. 1B), Greengenes (Fig. S1A) or GTDB (Fig. S1B). Moreover, because our amplicon library is linked to environments labeled with Earth Microbiome Project Ontology (EMPO) categories, we can easily identify the environments that contain samples that can fill out the tree. Because MAG assembly efforts can only cover abundant taxa, we plotted for each EMPO category the amount of new branch length added to the tree by taxa whose minimum abundance is 1% in each sample (Fig. 1C). The results show which environment types on average will best yield new metagenome assembled genomes (MAGs), and also show which environments harbor individual samples that will have a large impact when sequenced.

Past efforts to reconcile 16S and shotgun datasets have led to non-overlapping distributions and only techniques such as Procrustes analysis can even show relationships between the results ^14^. On two large human stool cohorts ^13,15^ where both 16S and shotgun data were generated on the same samples, we find that Bray-Curtis ^16^ (non-phylogenetic) ordination fails to reconcile at the feature level (Fig. 1D) and is poor at the genus level (Fig. 1E, S1C). However, UniFrac ^17^, a phylogenetic method, used with our Greengenes2 tree provides better concordance (Fig. 1F, S1D). To examine applicability of Greengenes2 to non-human environments, we next computed both Bray-Curtis and weighted UniFrac at the feature level on the 16S and shotgun data from the Earth Microbiome Project ^8^. As with the human data, we observe better concordance with the use of the Greengenes2 phylogeny (Fig. S2), despite limited representation of whole genomes from non-human sources as these environments are not as well characterized in general.

We also find that the per-sample shotgun and 16S taxonomy relative abundance profiles are concordant even to the species level. We first computed taxonomy profiles for shotgun data using the Woltka pipeline ^20^. Using a Naive Bayes classifier from q2-feature-classifier v2022.2 ^21^ to compare GTDB r207 taxonomy results at each level down to genus against SILVA v138 (Fig. 2A) or down to species against Greengenes v13_8 (Fig. 2B), no species-level reconciliation was possible. In contrast, Greengenes2 provided excellent concordance at the genus level (Pearson r=0.85) and good concordance at the species level (Pearson r=0.65) (Fig. 2C). Interestingly, the tree is now sufficiently complete that exact matching of 16S ASVs followed by reading the taxonomy off the tree performs even better than the Naive Bayes Classifier (Naive Bayes; Pearson r=0.54 at species, r=0.84 at genus).

**Figure 2.**
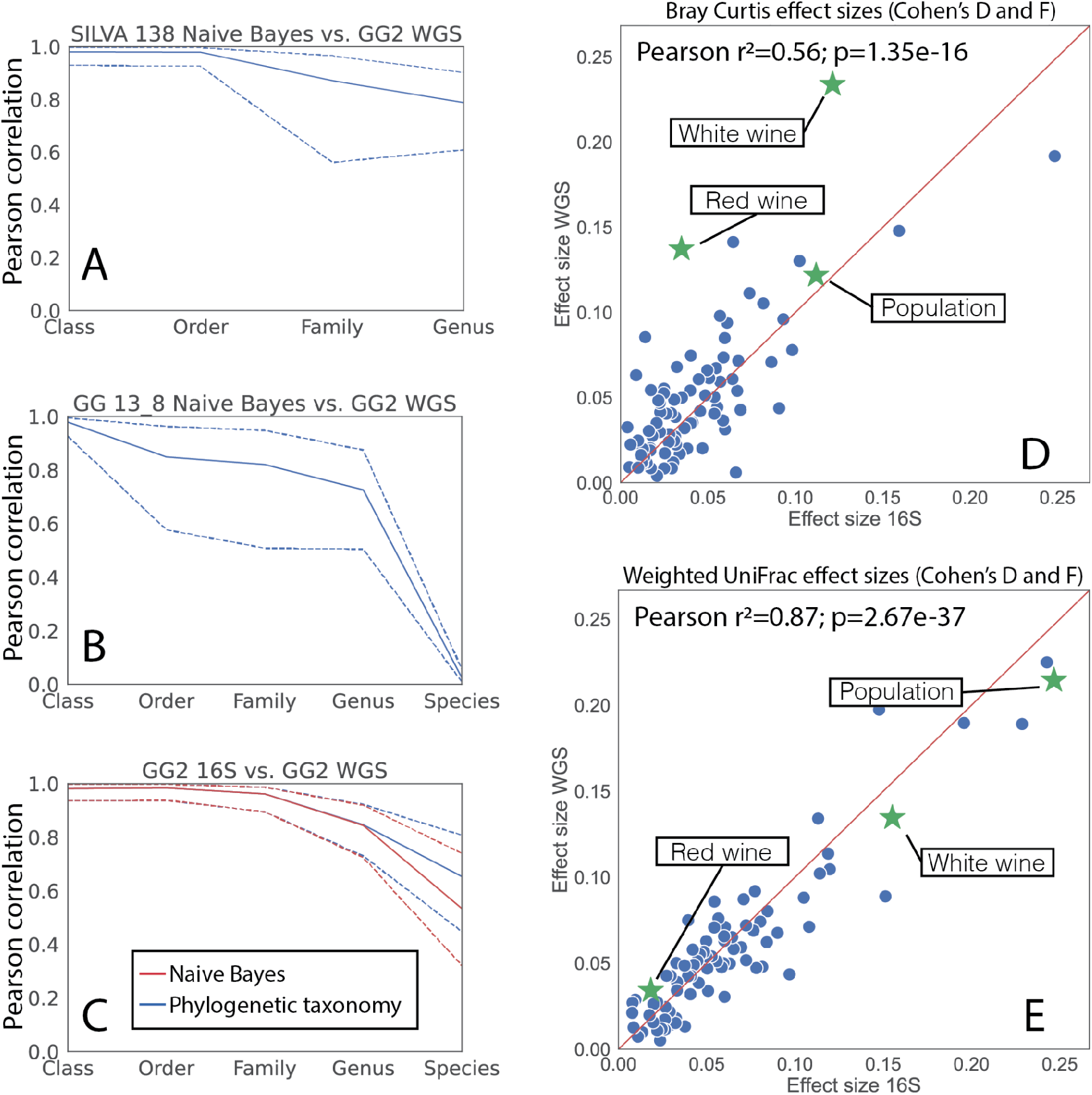
(A-C) Per sample taxonomy comparisons between 16S and whole genome shotgun profiles from THDMI. The solid bar depicts the 50th percentile, the dashed lines are 25th and 75th percentiles. (A) 16S taxonomy assessed with SILVA 138 using the default q2-feature-classifier Naive Bayes model (note: SILVA does not annotate at the species level). (B) 16S taxonomy assessment with Greengenes 13_8 using the default q2-feature-classifier Naïve Bayes model. (C) 16S taxonomy assessment performed by reading the lineages directly from the phylogeny or through Naive Bayes trained on the V4 regions of the Greengenes2 backbone. (D-E) Effect size calculations performed with Evident on paired 16S and whole genome shotgun samples from THDMI. Calculations performed at maximal resolution, using ASVs for 16S and genome identifiers for shotgun. The data represented here are human gut microbiome samples. The stars denote variables which are drawn out specifically in the plot (e.g., Population), and were arbitrarily selected as comparison points to help highlight differences between panels (D) and (E). (D) Bray-Curtis distances. (E) Weighted normalized UniFrac.

Finally, a critical reason to assign taxonomy is downstream use of biomarkers and indicator taxa. Microbiome science has been described as having a reproducibility crisis ^22^, but much of this problem stems from incompatible methods ^23^. We initially used the The Human Diet Microbiome Initiative (THDMI) dataset, which is a multipopulation expansion of The Microsetta Initiative ^13^ that contains samples with paired 16S and shotgun preparations, to test whether a harmonized resource would provide concordant rankings for the variables that affect the human microbiome similarly. Using Greengenes2, the concordance was good with Bray-Curtis (Fig. 2D; Pearson r^2^=0.56), better using UniFrac with different phylogenies (SILVA 138 and Greengenes2; Fig S1E; Pearson r^2^=0.77), and excellent with UniFrac on the same phylogeny (Fig. 2E; Pearson r^2^=0.87). We confirmed these results with an additional cohort ^15^ (Fig. S1FG). Intriguingly, the ranked effect sizes across different cohorts were concordant.

Taken together, these results show that use of a consistent, integrated taxonomic resource dramatically improves the reproducibility of microbiome studies using different data types, and allows variables of large versus small effect to be reliably recovered in different populations.

## ONLINE METHODS

### Phylogeny construction

Web of Life version 2 ^1^ (a tree inferred using genome-wide data) was used as the starting backbone. Full length 16S sequences from the Living Tree Project ^6^, full length mitochondria and chloroplast from SILVA 138 ^3^, full length 16S from GTDB r207 ^2^, full length 16S from Karst et al ^7^, and full length 16S from the EMP 500 ^8^ (samples selected and sequenced specifically for Greengenes2) were collected and deduplicated. Sequences were then aligned using UPP ^24^ and gappy sequences with less than 1000bp were removed. The resulting set of 321,210 unique sequences were used with uDance v1.1.0 to update the Web of Life 2 (WoL2) backbone. Briefly, uDance updates an existing tree with new sequences and (unlike placement methods) also infers the relationship of existing sequences. uDance has two modes: one that allows updates to the backbone and one that keeps the backbone fixed, where the former mode is intended for use with whole genomes. In our analyses, we kept the backbone tree (inferred using genomic data) fixed. To extend the genomic tree with 16S data, we identified 13,249 (out of 15,953 total) genomes in the WoL2 backbone tree with at least one 16S copy and used them to train a Deep-learning Enabled Phylogenetic Placement (DEPP) model with the weighted average method detailed below to handle multiple copies. We then used DEPP to insert all 16S copies of all genomes into the backbone and measured the distance between the genome position and the 16S position. We removed copies that were placed far further than others, as identified using a 2-means approach with centroids equals to at least 13 branches. We repeated this process in a second round. Then, for every remaining genome, we selected as its representative the copy with the minimum placement error and computing the consensus with ties. At the end, we were left with 12,344 unique 16S sequences across all the WoL2 genomes. For tree inference, uDance used IQ-TREE2 ^25^ in fast tree search with model GTR+Γ after removing duplicate sequences.

Next, we collected 16S V4 ASVs from Qiita ^10^ using redbiom ^26^ (query performed December 14, 2021) from contexts “Deblur_2021.09-Illumina-16S-V4-90nt-dd6875”, “Deblur_2021.09-Illumina-16S-V4-100nt-50b3a2”, “Deblur_2021.09-Illumina-16S-V4-125nt-92f954”, “Deblur_2021.09-Illumina-16S-V4-150nt-ac8c0b”, “Deblur_2021.09-Illumina-16S-V4-200nt-0b8b48”, “Deblur_2021.09-Illumina-16S-V4-250nt-8b2bff” and aligned them to the existing 16S alignment of sequences in WoL2 using UPP, setting the maximum alignment subset size to 200 (to help with scalability). The collected 16S V4 ASVs are aligned to the V4 region of the existing “backbone” alignments. A DEPP model was then trained on the full length 16S sequences from the backbone. DEPP constructs a Neural network model that embeds sequences in high dimensional spaces such that embedded points resemble the phylogeny in their distances.

Such a model then allows insertion of new sequences into a tree using distance-based phylogenetic insertion method APPLES-2 ^27^. The ASVs from redbiom were then inserted into the backbone using the trained DEPP model. To enable analyses of large datasets, we used a clustering approach with DEPP: we trained an ensemble of DEPP models corresponding to different parts of the tree and used a classifier to detect the correct subtree. During training, for species with multiple 16S, all the copies are mapped to the same leaf in the backbone tree. To train the DEPP models with multiple sequences mapped to a leaf, each site in the sequences is encoded as a probability vector of four nucleotides across all the copies.

### Integrating the GTDB and Living Tree Project taxonomies

GTDB and Living Tree Project are not directly compatible due to differences in their curation. As a result, it is not always possible to map a species from one resource to the other, either because parts of a species lineage are not present, are described using different names, or have an ambiguous association due to polyphyletic taxa in GTDB (e.g., Firmicutes_A, Firmicutes_B, etc, see https://gtdb.ecogenomic.org/faq#why-do-some-family-and-higher-rank-names-end-with-an-alphabetic-suffix for more detail). We integrated taxonomic data from LTP into GTDB as LTP includes species that are not yet represented in GTDB. Additionally, GTDB is actively curated, while LTP generally uses the NCBI taxonomy. To account for these differences, we first mapped any species that had a perfect species name association and revised its ancestral lineage to match GTDB. Next, we generated lineage rewrite rules using the GTDB record metadata. Specifically, we limited the metadata to records that are GTDB representatives and NCBI type material, and then defined a lineage renaming from the recorded NCBI taxonomy to the GTDB taxonomy. These rewrite rules were applied from most to least specific taxa, and through this mechanism we could revise much of the higher ranks of LTP. We then identified *incertae sedis* records in LTP that we could not map, removed their lineage strings and did not attempt to provide taxonomy for them, instead opting to rely on downstream taxonomy decoration to resolve their lineages. Next, any record that was ambiguous to map was split into a secondary taxonomy for use in backfilling in the downstream taxonomy decoration. Finally, we instrumented numerous consistency checks in the taxonomy through the process to capture inconsistent parents in the taxonomic hierarchy, consistent numbers of ranks in a lineage and ensuring the resulting taxonomy was a strict hierarchy.

### Taxonomy decoration

The original tax2tree algorithm was not well suited for a large volume of species level records in the backbone, as the algorithm requires an internal node to place a name. If two species are siblings, the tree would lack a node to contain the species label for both taxa. To account for this, we updated the algorithm to insert “placeholder” nodes with zero branch length as the parents of backbone records, which could accept these species labels. We further updated tax2tree to operate directly on .jplace data ^28^, preserving edge numbering of the original edges prior to adding “placeholder” nodes. To support LTP records which could not be integrated into GTDB, we instrumented a secondary taxonomy mode for tax2tree. Specifically, following the standard decoration, backfilling and name promotion procedures, we determine on a per record basis for the secondary taxonomy what portion of the lineage is missing, and place the missing labels on the placeholder node. We then issue a second round of name promotion using the existing tax2tree methods.

The actual taxonomy decoration occurs on the backbone tree, which contains only full length 16S records, and does not contain the amplicon sequence variants (ASV). This is done as ASV placements are independent, do not modify the backbone, and would substantially increase the computational resources required. After the backbone is decorated, fragment placements from DEPP are resolved using a multifurcation strategy using the balanced-parentheses library (https://github.com/biocore/improved-octo-waddle/).

### Phylogenetic collapse for visualization

We are unaware of phylogenetic visualization software that can display a tree with over 20,000,000 tips. To produce the visualizations in figure 1, we reduced the dimension of the tree by collapsing fragment multifurcations to single nodes, dropping the tree to 522,849 tips.

### MAG target environments

A feature table for the 27,015 16S rRNA V4 90nt Earth Microbiome Project samples was obtained from redbiom. The amplicon sequence variants (ASV) were filtered to the overlap of ASVs present in Greengenes2. Any feature with < 1% relative abundance within a sample was removed. The feature table was then rarefied to 1,000 sequences per sample. The amount of novel branch length was then computed, per sample, by summing the branch length of each ASV’s placement edge. The per sample branch length was then normalized by the total tree branch length (excluding length contributed by ASVs).

### Per sample taxonomy correlations

All comparisons used the THDMI^13^ 16S and Woltka processed shotgun data. These data were accessed from Qiita study 10317, and filtered the set of features that overlap with Greengenes2 using the QIIME 2^29^ q2-greengenes2 plugin. 16S taxonomy was assessed using either a traditional Naive Bayes classifier with q2-feature-classifier and default references from QIIME 2 2022.2, or by reading the lineage directly from the phylogeny. To help improve correlation between SILVA and Greengenes2, and Greengenes and Greengenes2, we stripped polyphyletic labelings from those data; we did not strip polyphyletic labels from the phylogenetic taxonomy comparison or the Greengenes2 16S vs. Greengenes2 WGS Naive Bayes comparison. Shotgun taxonomy was determined by the specific observed genome records.

Once the 16S taxonomy was assigned, those tables as well as the WGS Woltka WoL version 2 table were collapsed at the species, genus, family, order, and class levels. We then computed a minimum relative abundance per sample in the THDMI dataset. In each sample, we removed any feature, either 16S or WGS, below the per sample minimum (i.e., max(min(16S), min(WGS))), forming a common minimal basis for taxonomy comparison. Following filtering, Pearson correlation was computed per sample using SciPy ^30^. These correlations were aggregated per 16S taxonomy assignment method, and by each taxonomic rank. The 25th, 50th and 75th percentiles were then plotted with Matplotlib ^31^.

### Principal coordinates

THDMI Deblur 16S and Woltka processed shotgun sequencing data, against WoL version 2, were obtained from Qiita study 10317. Both feature tables were filtered against Greengenes2 2022.10, removing any feature not present in the tree. For the genus collapsed plot (figure 1e), both the 16S and WGS data features were collapsed using the same taxonomy. For all three figures, the 16S data were subsampled, with replacement, to 10,000 sequences per sample. The WGS data were subsampled, with replacement, to 1,000,000 sequences per sample. Bray-Curtis and Weighted UniFrac, and PCoA were computed using q2-diversity 2022.2. The resulting coordinates were visualized with q2-emperor ^32^.

The Earth Microbiome Project “EMP500” 16S and Woltka processed shotgun sequencing data, against WoL version 2, were obtained from Qiita study 13114. Both feature tables were filtered against Greengenes2 2022.10. The 16S data were subsampled, with replacement, to 1,000 sequences per sample. The WGS data were subsampled, with replacement, to 50,000 sequences per sample. The sequencing depth for WGS data was selected based on Figure S6 of Shaffer et al.^8^ which noted low levels of read recruitment to publicly available whole genomes. Bray-Curtis and Weighted UniFrac, and PCoA were computed using q2-diversity 2022.2. The resulting coordinates were visualized with q2-emperor.

### Effect size calculations

Similar to principal coordinates, the THDMI data were rarefied to 9,000 and 2,000,000 sequences per sample for 16S and WGS respectively. Bray-Curtis and weighted normalized UniFrac were computed on both sets of data. The variables for THDMI were subset to those with at least two category values having more than 50 samples. For UniFrac with SILVA, figure S1E, we performed fragment insertion using q2-fragment-insertion ^33^ into the standard QIIME 2 SILVA reference, followed by rarefaction to 9,000 sequences per sample, and then computed weighted normalized UniFrac.

For FINRISK, the data were rarefied to 1,000 and 500,000 sequences per sample for 16S and WGS, respectively. A different depth was used to account for the overall lower amount of sequencing data for FINRISK. As with THDMI, the variables selected were reduced to those with at least two category values having more than 50 samples. Support for computing paired effect sizes is part of the QIIME2 Greengenes2 plugin, q2-greengenes2, which performs effect size calculations using Evident (https://github.com/biocore/evident/).

### Data access

The official location of the Greengenes2 releases is http://ftp.microbio.me/greengenes_release/. The data are released under a BSD-3 clause license. A QIIME 2 plugin is available to facilitate use with the resource that can be obtained from https://github.com/biocore/q2-greengenes2/ (version 2023.3; DOI: 10.5281/zenodo.7758134). Taxonomy construction, decoration, and release processing is part of https://github.com/biocore/greengenes2 (version 2023.3; DOI: 10.5281/zenodo.7758138). uDance is available at GitHub: https://github.com/balabanmetin/uDance (version v1.1.0; DOI: 10.5281/zenodo.7758289).

Phylogeny insertion using DEPP is available at https://github.com/yueyujiang/DEPP (version 0.3; DOI: 10.5281/zenodo.7768798); the trained model accessioned with Zenodo at 10.5281/zenodo.7416684. The THDMI data are part of Qiita study 10317, and EBI accession PRJEB11419. The FINRISK data are available under EGAD00001007035. Finally, an interactive website to explore the Greengenes2 data is available at https://greengenes2.ucsd.edu.

## Acknowledgements

This work was supported in part by NSF XSEDE BIO210103, NSF RAPID 20385.09, NIH 1R35GM14272, NIH U19AG063744, NIH U24DK131617, NIH DP1-AT010885 and Emerald Foundation 3022. JTM was funded by the intramural research program of the Eunice Kennedy Shriver National Institute of Child Health and Human Development (NICHD). The Human Diets and Microbiome Initiative dataset was generated through support from Danone Nutricia Research and the Center for Microbiome Innovation.

---

**Figure S1.**
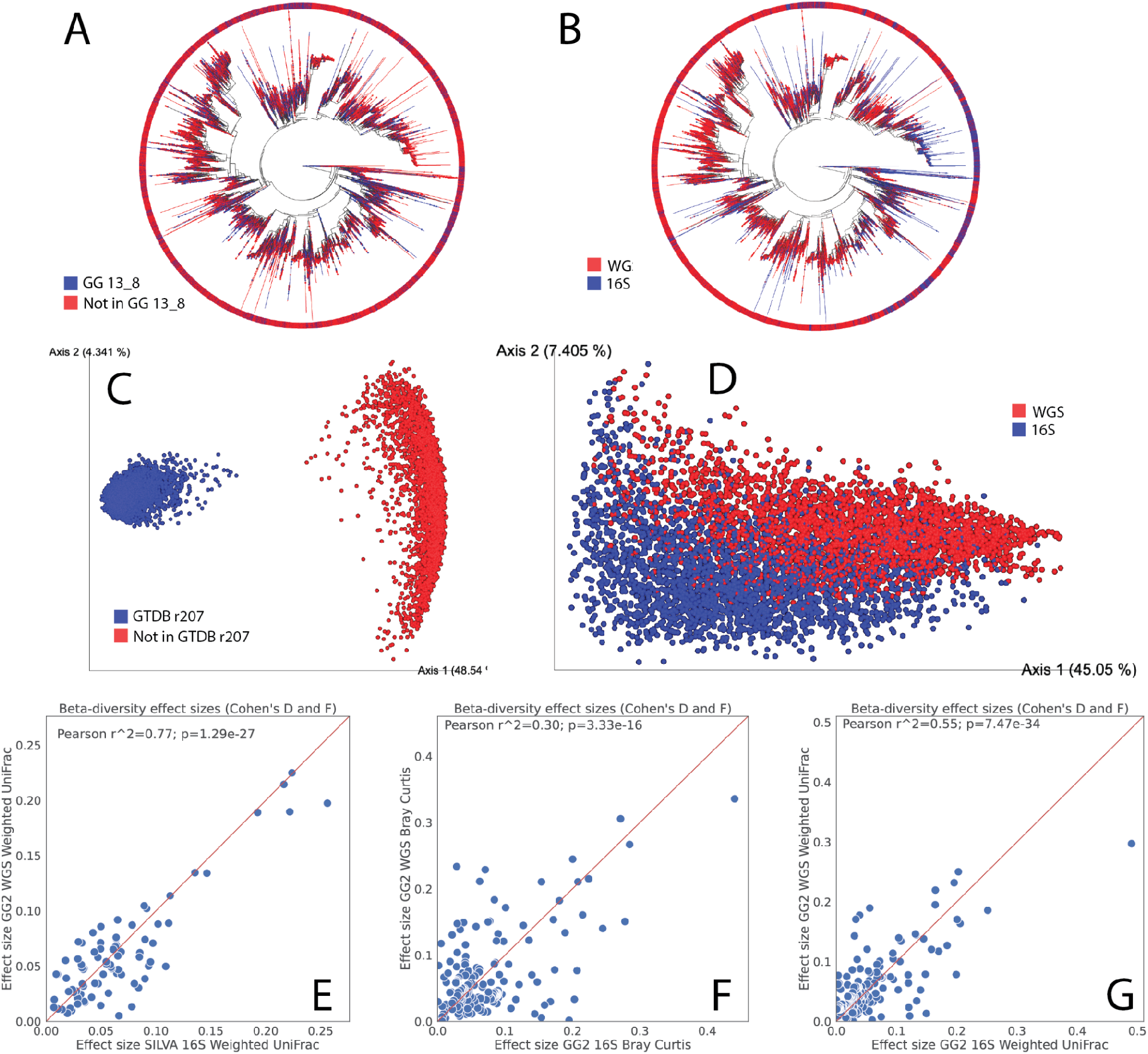
(A) Best BLAST hit for Greengenes 13_8 99% OTUs against Greengenes2. (B) Best BLAST hit for GTDB r207 SSU sequences against Greengenes2. (C) The FINRISK 16S and WGS data combined, collapsed to genus relative to Greengenes2, with Bray-Curtis computed followed by Principal Coordinates Analysis, colored by technical preparation. (D) The same data as (C) but using weighted UniFrac. (E) Effect sizes of the THDMI data using the SILVA 138 phylogeny for 16S data, and the Greengenes2 phylogeny for WGS data. (F) Effect sizes of the FINRISK data using Bray-Curtis. (G) The same data as (E) but using Weighted UniFrac.

**Figure S2.**
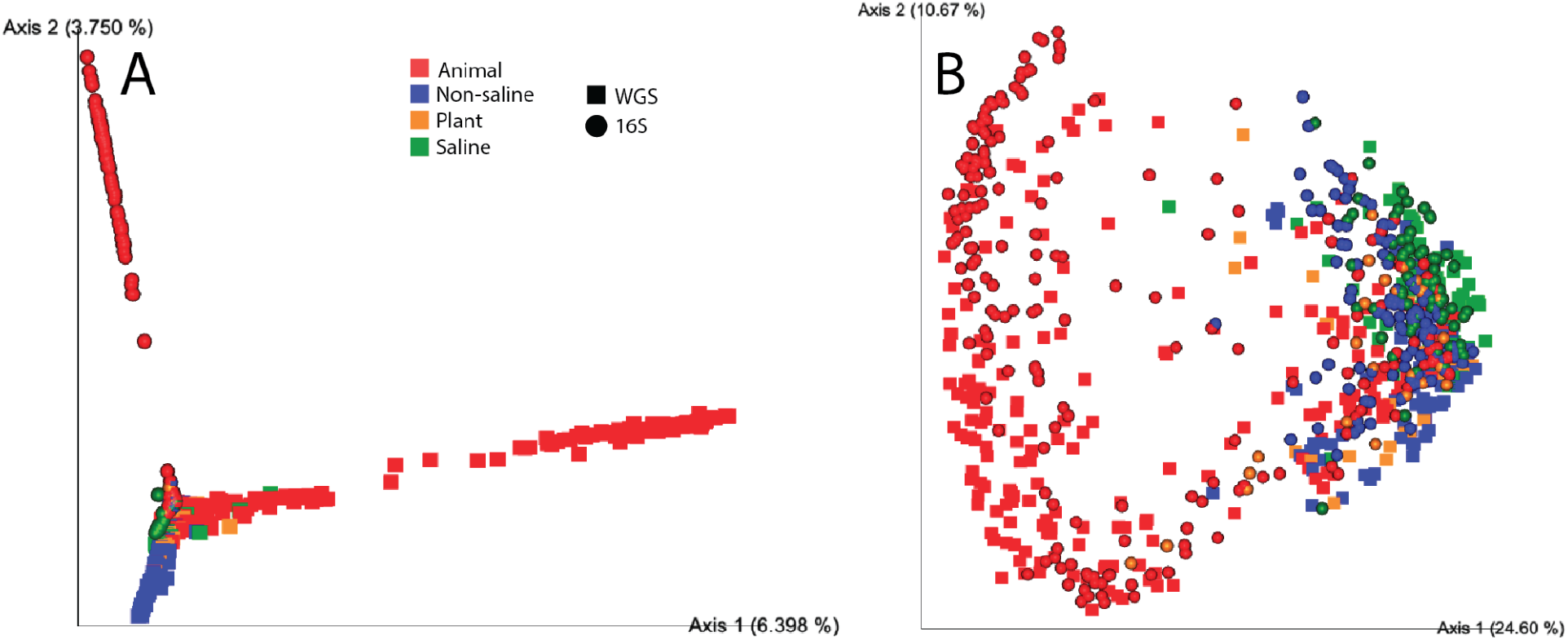
(A) The Earth Microbiome Project 16S and WGS data combined, and with Bray-Curtis computed at the feature level followed by Principal Coordinates Analysis, colored by Earth Microbiome Project Ontology level 2. Squares represent WGS data, and spheres represent 16S data. (B) The same data and coloring as in (A) but using weighted UniFrac.

